# Nucleosome conformational variability in solution and in interphase nuclei evidenced by cryo-electron miocroscopy of vitreous sections

**DOI:** 10.1101/295691

**Authors:** Mikhail Eltsov, Diana Grewe, Nicolas Lemercier, Achilleas Frangakis, Françoise Livolant, Amélie Leforestier

**Author notes:** Present address: IGBMC Imaging Center, CNRS, Inserm, Université de Strasbourg, 1 rue Laurent Fries, 67404 Illkirch, France.

## Abstract

In Eukaryotes, DNA is wound around the histone core octamer to form the basic chromatin unit, the nucleosome. Atomic resolution structures have been obtained from crystallography and single particle cryo-electron microscopy of identical engineered particles. But native nucleosomes are dynamical entities with diverse DNA sequence and histone content, and little is known about their conformational variability, especially in the cellular context. Using cryo-electron microscopy and tomography of vitreous sections we analyse the conformation of native nucleosomes, both *in vitro*, using purified particles solubilised at physiologically relevant concentrations (25-50 %), and *in situ*, within interphase nuclei. We visualise individual nucleosomes at a level of detail that allows us to analyse the conformation of the DNA wrapped around, and measure the distance between the DNA gyres. We evidence a variety of conformations. In interphase nuclei open nucleosomes predominate, with an average inter-gyre distance larger than that of the canonical particle. In concentrated solutions, we evidence a salt–dependant transition, with high salt compact conformations resembling the canonical nucleosome, and open low salt ones, closer to nuclear nucleosomes. Although further particle characterisation and cartography are needed to understand the relationship between this conformational variability and chromatin functional states, this work opens a route to chromatin exploration *in situ*.

## Introduction

The nucleosome is the basic packaging unit of eukaryotic chromatin and consists in 145-147 bp DNA wrapped into 1.65 turn of a left-handed superhelix around a histone octamer (two copies of H2A, H2B, H3 and H4). Atomic resolution structures are available since 1997 from X-ray crystallography (1-4) and more recently from cryoelectron microscopy (cryoEM) (5). All high resolution structures have been obtained from identical, symmetric and highly stable engineered particles, reassembled from recombinant histones and strongly-positioning DNA sequences. Together with the fact that nucleosomes are highly conserved structures, this has grounded a canonical view of the particle. This view is contrasted by the fact that the nucleosome is also a highly diverse particle, in terms of DNA sequence and protein content. Nucleosomes do not only play a role in DNA packaging, but are also involved in the regulation of many nuclear processes from gene expression to DNA replication and repair (6, 7). To allow and regulate chromatin functions, nucleosomes are thus not only modified (epigenetic modifications, histone post-translational modifications and DNA methylation), but also unwrapped, remodelled, disassembled, showing remarkable dynamical properties (8-12). Altogether, there is increasing evidence that there is no unique nucleosome structure but a whole family of conformations (13-15), little of which is still known. In addition, it is essentially documented *in vitro*. Within the nuclear context it is *terra incognita*.

Using cryo-electron microscopy of vitreous sections (CEMOVIS) we first analyse the conformation of nucleosomes in solution. We use native purified particles that reflect the chemical heterogeneity of nuclear nucleosomes, and analyse concentrated, physiologically relevant, solutions of isolated nucleosomes (25 to 50 % w:v). Under these conditions, nucleosomes are known to self-organise into multiple liquid crystalline (LC) phases depending on their concentration and ionic environment (16-20). In the presence of monovalent ions, an exhaustive phase diagram is available: above a concentration threshold nucleosomes pile up into columns, which in turn order the ones relative to the others to form different liquid crystals at higher concentrations (17). These phases constitute an easy to handle model system, with well defined and understood geometries and characteristics, which makes them ideal objects to explore the influence of many parameters (from ionic environment to nucleosome concentration), as well as sample preparation methods. We visualise nucleosomes in LC phases and analyse the conformation of the nucleosomal DNA. In particular, we access the distance *P* between the DNA gyres wrapped around the particle. We evidence variations of *P* and analyse these variations by comparison with the crystallographic structure (1, 2). This reveals conformational differences that correspond to an opening of the nucleosome in the direction of the superhelical axis. By comparing nucleosomes at different salt concentrations, we show that it is a salt-dependant phenomenon: low salt conformations are on average more open, while compact conformations closer to the canonical structure are found at higher salt.

The comparison of cryoEM data recorded on vitreous sections and thin films preparations of the LC phases shows that sectioning does not alter nucleosome conformation, making CEMOVIS a method of choice to access it not only *in vitro* in concentrated solutions, but also in its native nuclear context.

We thus question the occurrence of open conformations *in situ*, investigating nucleosomal DNA in interphase cell nuclei from different cell types, two cultured human cell lines, and Drosophila embryos. We obtain both 2D images and cryo-tomograms of vitreous sections where nucleosomes can be recognised individually and analysed at a high enough level of detail to visualise DNA wrapped around the particle and measure *P*. On average, nucleosomes *in situ* are more open than the canonical crystallographic structure, with *P*-value closer to that of native purified particles in low salt solution. Yet the particle’s variability is more important, probably reflecting the different functional states found in the cell nucleus.

## Material & Methods

Nucleosome purification and preparation of concentrated solutions. Native H1-depleted nucleosomes (histone octamer associated with 167 ± 10 bp DNA) and native nucleosome core particles (146 ± 3 bp DNA) were respectively prepared from calf thymus and chicken erythrocyte chromatin. Briefly, chromatin was extracted in low ionic strength buffer, followed by histone H1 removal. Nucleosome particles with different DNA associated lengths were obtained by controlled digestion with micrococcal nuclease, and purified by gel filtration (Sephacryl S300 HR, GE Healthcare). The integrity of all core histones, and the absence of contaminating di- and oligo-nucleosomes were checked by gel electrophoresis. Purified particles were extensively dialyzed at a concentration of 1 mg/ml against TE buffer (10 mM Tris pH 7.6, 1 mM EDTA) supplemented with either 5 mM NaCl (low salt) or 150 mM (high salt). The solutions were concentrated by ultrafiltration through a cellulose membrane (Ultracell YM100, Millipore) in a pressurized cell up to about 250 mg/ml (25% w:v). Higher concentrations were reached by osmotic pressure, induced either by extensive dialysis against PolyEthyleneGlycol (PEG, MW20000) dissolved in the same buffer at concentrations ranging from 10 to 22 % (w:v), or by progressive mixing with the PEG solution, as described in (20).

### Cell culture and preparation

Cell suspensions of the human cell lines HT29 (21) and KE37 (22) were pelleted by centrifugation (3000g, 3 min), and gently re-suspended in an equal volume of dextran (PM 40000, Sigma Aldrich) dissolved at a concentration of 40% (w:w) in Phosphate-Buffered Saline (PBS). *Drosophila melanogaster* (Bloomington Stock number 30564) were maintained on standard Bloomington medium. Embryos were collected at apple agar plates, dechorionated in 50% (v:v) bleach (Sigma-Aldrich) and washed with distilled water. Developmental stages 14-15 were selected by visual inspection under a stereomicroscope (Leica M165 FC) and transferred into gold-plated copper carriers with a 0.1 mm indentation (B-type, Engineering Office M. Wohlwend GmbH) filled with 25% dextran in PBS and processed for high-pressure freezing.

### Bulk sample vitrification and cryo-ultramicrotomy

Nucleosome LC solutions and cell pellets were frozen by slam-freezing against a copper block cooled down to liquid helium temperature (Cryovacublock, Reichert) as described in (23). *Drosophila* embryos were frozen by high pressure freezing using HPM 010 machine (ABRA Fluid AG). Frozen specimens were then transferred into a Leica FC6/UC6 ultramicrotome (Leica Microsystems) installed in a controlled environment with a relative humidity maintained below 20%. Nucleosome solutions and cell suspensions were sectioned at −160 °C, Drosophila embryos at −145 °C, using a 25° or a 35° diamond knife (Diatome). The cutting feed was set at 40 nm for sections dedicated to 2D imaging, in order to minimize superimposition effects. A cutting feed of 75 nm was used for thicker sections dedicated to tomography. Sections were respectively collected on Quantifoil S7/2 grids coated by a continuous thin carbon film (2 D imaging), or directly on C-flat CF-2/1 grids (Electron Microscopy Sciences) for tomography. They were pressed against the grid using a mechanical press (24) or an electrostatic press (Crion (25), Leica).

### Thin film vitrification

Drops of LC nucleosome solutions were deposited onto an electron microscopy grid covered with a holy carbon film. On account of their high viscosity, drops were strongly blotted manually between two filter papers, and immediately vitrified by immersion in liquid ethane using a home-made plunging device. To avoid water evaporation, the procedure was performed under humidified atmosphere (26).

### Cryoelectron microscopy

For 2D imaging, grids were mounted in a Gatan 626 cryoholder (Gatan, Pleasanton, USA) and transferred in a JEOL 2010 Field Emission Gun (FEG) transmission electron microscope (TEM) operated at 200 kV or a FEI CM12 TEM operated at 80 kV. The vitreous state of the specimens was checked by electron diffraction. Images were recorded on Kodak SO163 films, under low-dose conditions at direct magnification of 25,000× to 45,000×, with defocus in the range of 1.5 μm (200kV) and 900 nm (80 kV). Films were developed in full-strength Kodak D19 for 12 min. In these imaging conditions, the signal of interest (typically in the range of 2.5 to 3.5 nm, lies far below the first zero of the contrast transfer function (CTF) (1.9 nm for both imaging conditions). We checked that CTF correction is not necessary for image interpretation (see Supplementary Figure S1). Images presented in Fig. 1-3 were denoised by wavelet filtration using the ImageJ “A trous wavelet filter” plugin (with k_1_ = 20, k_2_ = k_3_ = k_4_ = k_5_ = 0).

**Figure 1.**
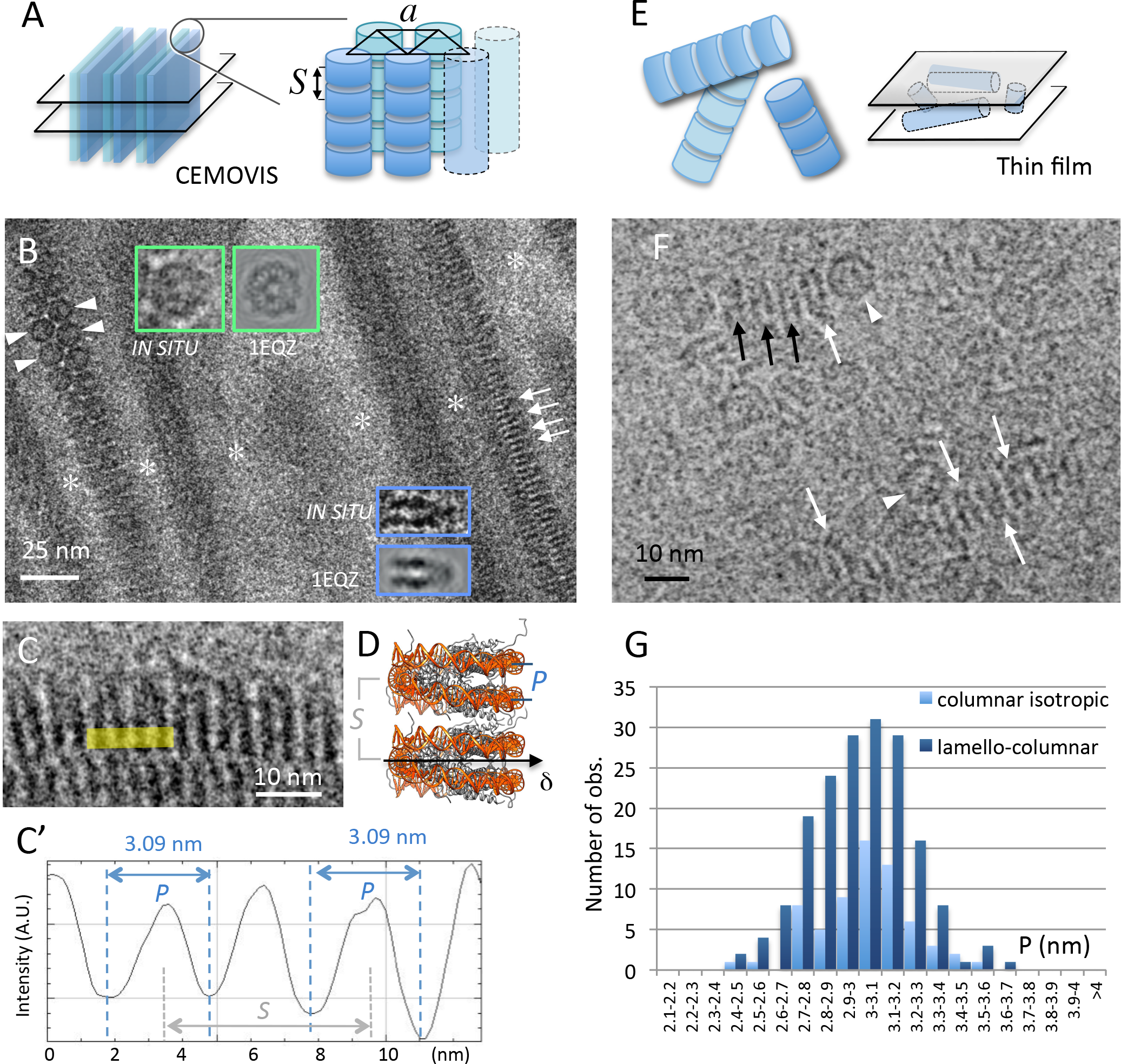
CryoEM of nucleosomes particles forming columnar phases in low salt solutions (15 mM [+]). (A) The lamello-columnar phase is composed of piles of nucleosomes (columns) that align in parallel onto a triangular lattice of parameter *a*, forming bilayers. Bulk phase (long range ordered bilayers) can be sectioned (eg transversally, left) and analysed by CEMOVIS. (B, C) CEMOVIS imaging of the lamello-columnar phase. Bilayers in transverse section are separated by particle-free layers of solvent (white stars). Piles of nucleosomes are seen in top (arrowheads), side (arrows) or oblique views. Top and side views are compared to the corresponding projections of the crystallographic structure of the NCP (from PDB 1EQZ) in the inserts. (C, C’) Measurement of P values on columns in side view. (D) In the columns of the lamello-columnar phase, nucleosomes stack on top of each other with parallel alignment of their dyad axes (δ). P corresponds to the distance between the DNA gyres measured at the front side of the particle. S corresponds to the stacking repeat. (F) In the columnar isotropic phase observed by thin film cryoEM piles of nucleosomes form a random network sketched in (E). Nucleosomes are recognized in side (arrows) and top (arrowheads) views. (G) Distribution of *P* values measured in the lamello-columnar and columnar isotropic phases. (B, F) White arrows indicate the orientation of the nucleosome’s dyad axis δ where it lies in the observation plane. Black arrows point to side view particles with their dyad axis out of the observation plane.

For cryoelectron tomography were mounted into Autogrid rings (FEI, Thermofisher, Eindhoven, The Netherlands) and transferred into a Titan Krios (FEI, Thermofischer, Eindhoven, The Netherlands) operated at 300 kV equipped with GATAN GIF Quantum SE post-column energy filter and K2 Summit direct electron detector (Gatan, Pleasanton, USA). Tilt series were recorded using Serial EM software (bio3d.colorado.edu/SerialEM (27)) at a nominal magnification of 64000× (2.2 Å/pixel). The dose-symmetric recording scheme macro (28), was run within an angular range from – 60° to + 60°, with a starting angle 0° and an angular increment of 2°. The electron dose was set to 1.5 e–/A^2^ for individual tilt images, corresponding to the total dose 91.5 e–/A^2^ for the complete tilt series. Contrast of tilt images recorded with imaging conditions used for 2D images (CTF first zero at 1.9 nm) was not sufficient for fiducial-free alignment. A reliable alignment was achieved when the target defocus was set to − 3.5 µm, corresponding to the CTF first zero at 2.4 nm. Since this value is relatively close to the signal of interest (*P* sampling range, see Results), CTF correction was performed for all tomographic data. Generation of fiducial-free markers was done by patch tracking algorithm of IMOD package (29). Tilt series alignment based on marker positions generated by IMOD, CTF correction and tomogram reconstruction were performed with customs scripts (30,31). The reconstructed volumes were denoised by filtering by 3D nonlinear anisotropic diffusion (k=1, 10 iterations) (32). 5 nm tomographic slices through the reconstructed section volumes were generated in IMOD.

### Generation of CTF-modulated projections and tomographic slices from the canonical NCP

For simulation of 2D nucleosome reference images, a 3D electron density map was generated from 1EQZ PDB map using pdb2em function of EMAN software package (33) with a resolution of 1 nm and pixel size 0.025 nm. All following calculations were done with using custom scripts running under Matlab platform (MathWorks, Natick, USA). The nucleosome electron density map was rotated in space with all three rotational angles assigned to random numbers, and a projection of the complete volume was generated. CTF was applied to the projection image with defocus assigned as a random number within the defocus range of real 2D images of nucleosomes acquired at 200 kV (-1.5 ± 0.5 μm) and at 80 kV (900 nm). This procedure was repeated to generate a set of reference images (Supplementary Figure S2). Similar patterns were obtained for both imaging conditions. To simulate 3D reference images, the 1EQZ nucleosome electron density map was binned down to 64×64×64 pixel volumes with the final voxel of 0.2 nm. We generated a 75nm thick virtual chromatin section, as a volume of 1024×1024×375 pixels where 750 randomly oriented nucleosomes were placed into random positions, without density overlapping. A series of tilt images was generated by rotation and projection of each volume, using the same angle range increment as for real tomograms. We found that, in our tomograms of vitreous sections, an average defocus corresponded to -3.7 µm at 0° tilt, while it fluctuated from -3 µm to -4.5 µm at higher angles. Accordingly, 0° tilt images of the generated volumes changes were modulated with CTF for the defocus of -3.7 µm, whereas defocus values for other tilt angles were generated as a list of random numbers within the aforementioned range. The generated tilt series were then aligned, CTF-corrected and reconstructed with the same scripts as used for the real tomogram reconstructions. Similarly, 5 nm thick tomographic slices were generated in IMOD.

### P-measurements and statistical analyses

Line profiles measurements were performed using the ImageJ software (34). Shapiro-Wilk (SW) test was used to assess normality of *P* distance measurements distributions. All data samples were distributed normally with the exception of High-Salt solutions measurements. The two-sample t-test was used to assess difference between normally distributed data samples. The Kolmogorov-Smirnov test was used to assess difference between High Salt and other data samples. All test were performed with alpha level of 0.05. Statistical analysis was done using XLSTAT software (Addinsoft, New York, USA).

## RESULTS

### CryoEM of nucleosomes in concentrated solutions

LC phases of nucleosomes were prepared at two different concentrations of added monovalent salt: 15 mM (low salt) and 160 mM (high salt), in order to question the effect of salt concentration on nucleosome conformation. We analysed both H1-depleted nucleosomes and nucleosome core particles (DNA lengths of 167±10 and 145±3 bp respectively, see Material & Methods). Within the range of concentration explored here (25-50 % w:v), we found no difference, both in terms of LC organisation and particle conformation, between the two sets of particles. They will hereafter simply be referred as “nucleosomes core particles” (NCP).

#### Low salt solutions: evidence for a variable nucleosome conformation, on average more open than the canonical crystallographic structure

At low added salt concentration (≤ 40 mM), and at NCP concentrations between 28 and 32 % (w:v), columns of NCP order with a local triangular lattice, into a lamello-columnar phase (Figure 1A). This phase was previously analysed by CEMOVIS, providing a molecular scale understanding of nucleosome order and interactions (20). Figure 1B, shows a transverse section of the lamello-columnar phase, with bilayers of columns in apical and side views. In both cases, the orientation of the NCP can be determined by comparison with an image gallery generated from the PDB (1EQZ), applying the contrast transfer function (CTF) corresponding to our imaging conditions. The overall shape of the NCP is that of a flat cylinder, about 11.5 nm in width and 6 nm in height, with DNA wrapped into 1.65 turn on its lateral surface. Multiple patterns are expected (Supplementary Figure S2). Both disc-shaped top views, and highly contrasted striated side views, characterised by a pair of stripes drawn by the nearly two gyres of DNA wrapped around the particle, are unambiguously identified (Figure 1B inserts and pointed by arrowheads and arrows respectively). Top views reveal the triangular lattice formed by columns. Side views present long-range periodic striations corresponding to DNA of piled particles. The period *S* corresponds to the stacking repeat of the NCPs (Figure 1C, sketched in 1D). The distance *P* characterises the distance between the two gyres of DNA wrapped around a given particle. We systematically measured *P* on intensity line profiles using ImageJ (Figure 1C’), wherever NCP are oriented in side views. The *P* distribution is given in Figure 1G (dark blue): it varies from 2.5 to 3.67 nm, and follows a normal distribution (Shapiro-Wilk (SW) test p = 0.399), with an average value of 2.99 ± 0.23 nm.

##### CEMOVIS-associated compression does not affect nucleosome structure

The major artefact associated to the CEMOVIS approach is the compression induced during the cutting process (35-37). Compression occurs in the cutting direction *k* and is coupled to an increase of the section thickness. It has been reported to induce severe deformations, up to 20-30 %, of long-range structures e.g. entire bacterial or yeast cells, or organelles. Supramolecular assemblies, such as microtubules, were also reported to deform. In contrast, at the molecular level, deformations have been shown to be negligible or even inexistent: the structure of ribosomes, as well as the local packing of DNA in sperm cells or crystalline protein lattices were shown to be un- or little affected (38-40). To determine whether or not compression modifies the *P* parameter of the NCP, we sorted out a subset of measurements obtained on the lamello-columnar phase *in vitro.* It appears that most of the *P* values were recorded on side view NCP oriented between 0 and 45° to the cutting direction *k.* Only few were observed oriented perpendicular, suggesting that long-range compression may perturb chromatin visualisation, possibly inducing local reorientations. Nonetheless, where clearly visible side view NCP were oriented perpendicular to k, *P* did not show any variation. We also compared *P* for particles oriented at 0(±5)°, 45(±5)° and 90(±5)° to *k*, and this analysis confirms that there is no compression effect on the *P* value (Supplementary Figure S3).

##### CEMOVIS *versus* thin film cryoEM: the sample preparation procedure does not influence nucleosome structure

Single bilayers of columns can also be trapped within thin films (Supplementary Figure S4). Bilayers are then always parallel to the observation plane, and the columns of the two layers superimpose within the film thickness. Side views of individual nucleosomes are recognised in some places, and *P* values measured on line profiles as described above. *P* also follows a normal distribution (SW p-value 0.863), and varies from 2.41 to 3.51 nm (3 ± 0.21 nm). The comparison with CEMOVIS measurements (Student test p-value 0.519; see Fig. S4) shows that there is here no influence of the sample preparation method.

##### The inter-distance and relative orientation of stacked particles do not influence the nucleosome conformation

The stacking distance *S* between particles varies with the particle concentration, from 5.9 to 6.2 nm in the lamello-columnar phase (41), and from

5.9 to about 7 nm within the less concentrated columnar isotropic phase (25-28 % w:v). The local order between neighbouring particles also differs in both phases (20). NCP always pile in parallel in the lamello-columnar phase (with their dyad axis oriented inward the bilayer, Figure 1B, white arrows and arrowheads). In contrast, there is no preferred orientation of NCP dyad axes within columns of the isotropic phase, where parallel as well as anti-parallel and intermediate orientations are found. This is observed in Figure 1F, which shows a local view of the columnar isotropic phase trapped in a thin film. *P* varies from 2.48 to 3.51 nm in the columnar isotropic phase (3.03 ± 0.2 nm). The *P* distribution is normal (Shapiro-Wilk 0.606) and can be compared to that found for the lamello-columnar phase (Figure 1G). No significant difference can be evidenced (t-test p-value 0.197), which suggests that both the local stacking density and the relative orientation of neighbours do not significantly influence the conformation of stacked NCP.

NCP in low salt columnar solutions present a range of conformations. The global set of *P* data obtained from the different low salt phases follows a normal distribution (SW p-value 0.551), with an average value of 3 ± 0.22 nm. In order to compare these data with the canonical crystallographic structure, we measured *P* on the different striated projection patterns generated from the PDB file (Supplementary Figure S2, S5), exactly as it was done on EM images. This also results in a normal distribution (SW p-value 0.142), but with an average value of 2.79 ± 0.14 nm (Supplementary Figure S5), which is significantly different from *P* in low salt solutions (t-test p-value 9.4 10^−15^). This comparison reveals that native NCP in low salt solution markedly differ from the particle’s canonical PDB structure: they present a whole range of conformations rather than a single one, and are on average more open than synthetic particles in crystals.

#### High salt solutions: nucleosomes adopt more compact conformations resembling the canonical structure

Above 100 mM added salt concentration, dense columnar nematic and columnar hexagonal phases form, at NCP concentrations higher than 45% and 50% (w:v) respectively (17, 18, 41). Both vitreous thin films (Figure 2A) and sections (not shown) were obtained from hereafter so-called “high salt” (160 mM) columnar solutions, and imaged by cryoEM. In both phases, striated patterns typical of stacked side view particles are observed. In thin films, columns are oriented parallel to the film and superimpose; a local nematic order can be detected locally (Figure 2B). Individual NCP are recognised and *P* measured on line profiles (Figure 2C). Measurements obtained from thin film cryoEM (mean 2.81 nm, s.d. 0.218 nm) and CEMOVIS (mean 2.78 nm, s.d.0.22 nm) do not show a significant difference (Kolmogorov-Smirnov test p = 0.588, α = 0.05). They were therefore combined into a single data sample, whose distribution is given in Figure 2D (in green), and compared with low salt data (in blue). The Shapiro-Wilk test (with a p-value equal to 0.032) does not allow us to hypothesize a normal distribution for *P* at high salt. To compare the high salt and low salt populations, we therefore use a Komolgorov-Smirnov test, resulting in a p-value smaller than 0.0001 (α = 0.05). While in both cases *P* presents a broad distribution, it is larger in low salt than in high salt solutions. We also compared the high salt data with the simulated measurements from PDB. The mean *P* value at high salt (2.79 nm) is the same as that simulated from PDB, thus supporting a similarity between native NCP in high salt solution and synthetic particles in crystals. Nevertheless the distribution of the high salt data is significantly different from the simulated measurements (Komolgorov-Smirnov test, p-value = 0.022; α = 0.05). It is broader (s.d. 0.23 versus 0.14 nm), and enriched in larger *P*-distances, typical of the low salt particles.

**Figure 2.**
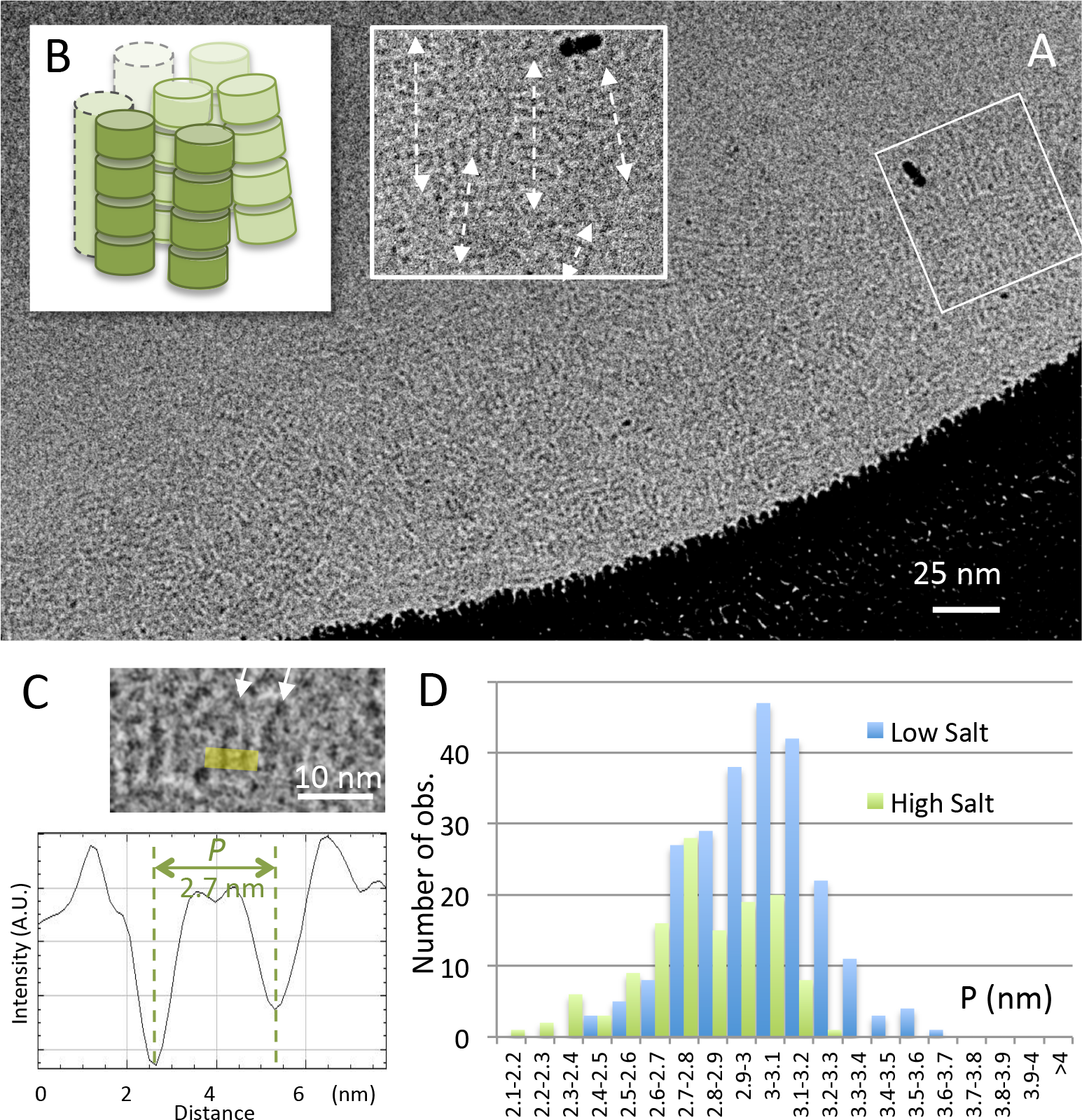
Comparison of *P* values measured in solutions of purified nucleosome solutions under low *versus* high salt conditions. Columnar phase formed in high salt (160 mM [+]) (A) trapped within a thin film and observed by cryoEM. Columns align parallel to the film and superimpose within its thickness, leading to complex striated patterns. (B) Observation of a local nematic order in an enlargement of A (white square).(C) Side view nucleosomes are recognized locally and *P* is measured on line profiles. (D) Distributions of *P* values recorded in the columnar hexagonal phase (160mM [+], green) and in the lamella-columnar phase (15 mM [+], blue).

Altogether, these observations reveal that nucleosome undergo a salt dependant conformational change at physiologically relevant nucleosome concentrations. The canonical particle resembles native nucleosomes in high salt solutions, while more open conformations are found in low salt environment.

By comparing thin film cryoEM and CEMOVIS approaches, we demonstrate that vitreous cryo-sectioning does not alter the NCP conformation. Note that, although it is not the focus of the present paper, CEMOVIS approaches also do not disturb significantly inter-particle interactions and ordering, as already shown *in vitro* (17). Having thus shown that CEMOVIS is well adapted to access nucleosome conformation in crowded environments, we carried out a CEMOVIS analysis of nucleosomes in their nuclear context. We prepared vitreous sections of interphase nuclei of different cell types, two human cell lines (HT29 and KE 37) and *Drosophila melanogaster* embryonic brain cells. HT29 and KE37 were imaged in 2D, and cryo-tomograms were recorded from *Drosophila* embryos.

### Cryo-electron microscopy and tomography of vitreous sections (CEMOVIS and CETOVIS) of interphase chromatin

#### Nucleosomes can be visualised and their conformation analysed in the interphase nucleus

Figure 3A shows a local view of the nucleus of a KE37 cell, with its nuclear envelope (NE) and a nuclear pore complex (NPC). Chromatin presents a homogeneous grainy texture resembling that already described (42-44). Yet, this texture, observed in the two cell lines, is here finer than classically reported in CEMOVIS studies, on account of our choice of lower underfocus imaging. Dense chromatin rich regions are interspaced by less dense regions, yet with no well defined frontiers (Figure 3B). As already described in mammal cells, dense chromatin regions are always found in the vicinity of nuclear envelope (45). In addition, we observe striated patterns (Figure 3B, C, arrows). These striations are ubiquitous in the nucleus, and are found both in dense and less dense regions. They may remain very local, corresponding to a single pair of stripes (Figure 3B, C, black arrows), or propagate over larger distances (Figure 3B, C, white arrows) either in the streak direction (up to 60-100 nm) or perpendicularly (typically 10-15 nm and up to 40-45 nm in a few cases).

**Figure 3.**
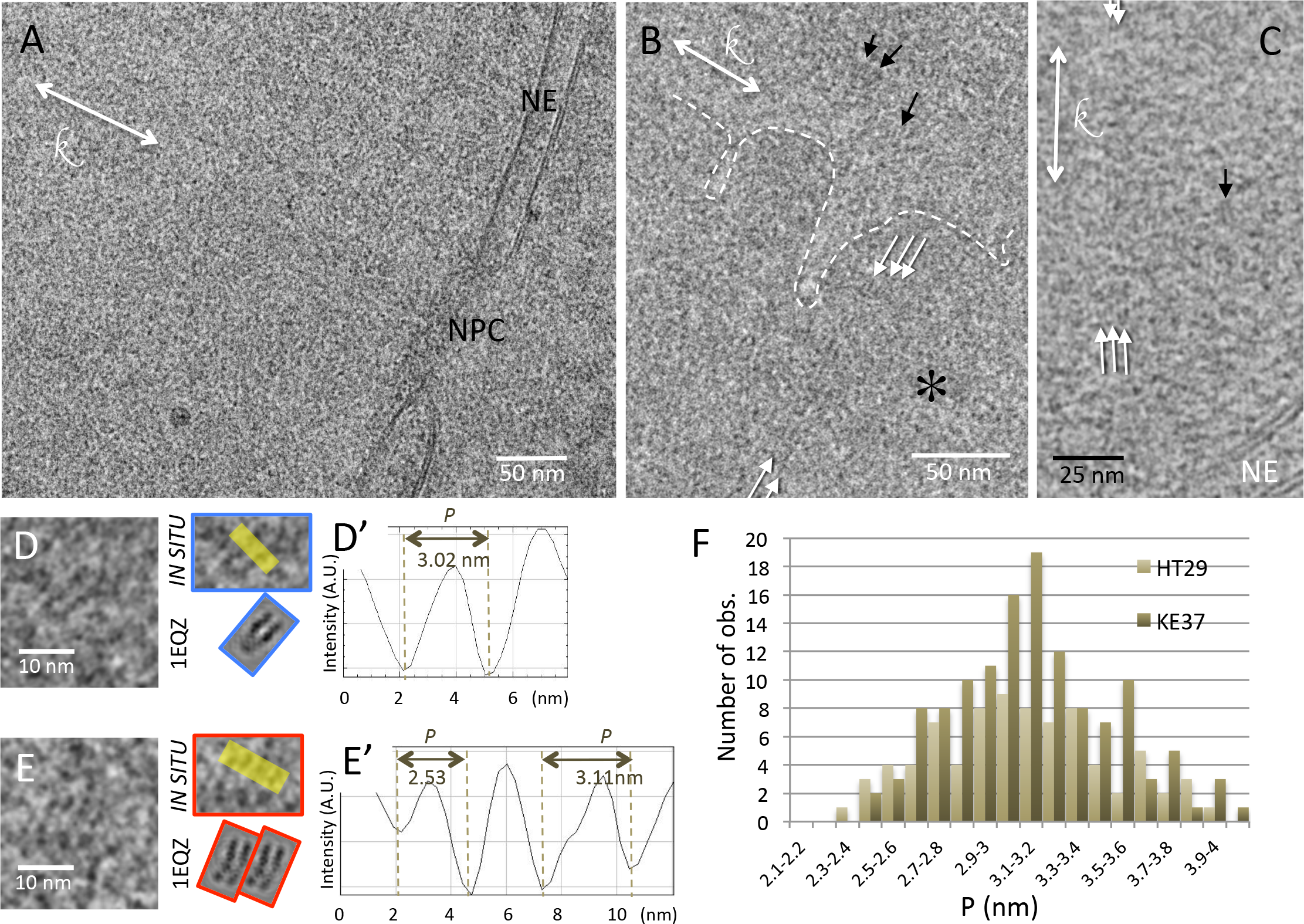
CEMOVIS of interphase nuclei of the human cell lines KE37 (A, B, D) and HT29 (C, E). (A) Local view of a nucleus limited by the nuclear envelope (NE). A nuclear pore complex (NPC) is visible. (B) A chromatin-dense region (*), delineated by the white dotted line, coexists with a less dense region. (B, C) Striations are visible, either locally (black arrows) or extending over longer distances (white arrows). (D, E) Nucleosomes visualised in side views. The patterns are compared with CTF-modulated projections of the crystallographic structure of the particle (from PDB 1 EQZ). (D’, E’) The distance between the DNA gyres *P*, is measured on line profiles indicated in yellow in (D) and (E).(F) Distribution of *P* for the two cell types.

Interphase chromatin is a complex and heterogeneous structure. 2D projection images are therefore far more complex to interpret and analyse than those obtained with ordered LC *in vitro*, where NCP are the only macromolecular species and pile up in register. Yet, nucleosomes are the major chromatin component. By comparison with the gallery of projections (Supplementary Figure S2) and images of NCP in solution (Figure 1, 2), individual side view nucleosomes are clearly recognised and their orientation can be determined (Figure 3D, E). In contrast, apical and oblique views could not be unambiguously recognised. This can be understood by a combination of two phenomena that favour the identification of side views. In all cases, within the thickness of a section, a few nucleosomes, possibly variously oriented, are expected to superimpose, leading to intricate patterns. Pattern analysis can also be more complex due to the presence of a number of other macromolecular species whose nature and amount varies with space and time. A single nucleosome oriented perpendicular to the section plane (side view) occupies about a fourth of the section thickness, *versus* a sixth when it lies parallel to the section (apical view). Superimposition effects are therefore less severe in the first case. In addition, the typical striated patterns are about twice more contrasted than apical and oblique views (Supplementary Figure S2). The signal of a side view nucleosome can therefore be expected to dominate other contributions, which facilitates its identification.

To avoid superimposition effects of 2D imaging and access the conformation of variously oriented nucleosomes, we recorded cryo-tomograms of vitreous sections. We found that acquisition of tilt series is much more demanding; therefore obtaining the highest density of nuclei within the section is essential to find locations suitable for tomography. Since organ/tissue samples better fulfil this criterion than cell suspensions, we chose the developing *Drosophila* brain. It shows both a high cell density and high nuclei/cytoplasm ratio, contributing to the yield of suitable chromatin areas (46). In addition, *Drosophila* embryos can be efficiently vitrified (47) and are surrounded by the liquid-impermeable vitelline membrane (48,49), which remarkably preserves the native extracellular environment during sample preparation and freezing.

Chromatin of embryonic *Drosophila* brain cells is organized into two main regions: the highly-compact chromocenters which accumulate constitutive heterochromatin regions of all chromosomes attached to nuclear envelope (not shown), and less compact domains containing facultative heterochromatin and euchromatin. These are distributed within nucleoplasm and proved more accessible for cryo-tomography analyses. Figure 4A shows a typical 5-nm thick section extracted from a tomogram. The nuclear envelope with its nuclear pore complex (NPC) is recognised. Within the nucleus, disperse chromatin-dense regions are identified (Figure 4A, dotted lines). As in 2D images, striations are locally observed (arrows). Chromatin rich regions are surrounded by the low density nucleoplasm. In Figure 4B, C, the local observed patterns are compared with the gallery of 5-nm thick projections generated from PDB maps of the NCP (1EQZ) applying the CTF-modulation and correction of the tomograms (Supplementary Figure S6). Our analysis unambiguously reveals not only striated patterns, which are representative of side views (and are the only recognizable ones in 2D images), but also all other characteristic patterns, including top views (Figure 4B, green). Amongst the variety of side views, it is now possible to fully determine the particle orientation (as sketched in Figure 4D), and in particular to distinguish back and front views when the nucleosome dyad axis lies perpendicular to the section plane. Figure 4C shows examples of back (red) and front (yellow) views showing the entry/exit DNA sites. We checked the compatibility of the xy view of each particle, with its xz and yz views (Figure 4E), which unambiguously proves that the observed patterns do indeed correspond to a nucleosome. In addition, these 3D data allowed us, in a few cases, to determine the handedness of the DNA superhelix wrapped around the histone octamer (Supplementary Figure. S7). 12 particles were analysed, all showing a left-handed superhelix, as in the canonical structure.

**Figure 4.**
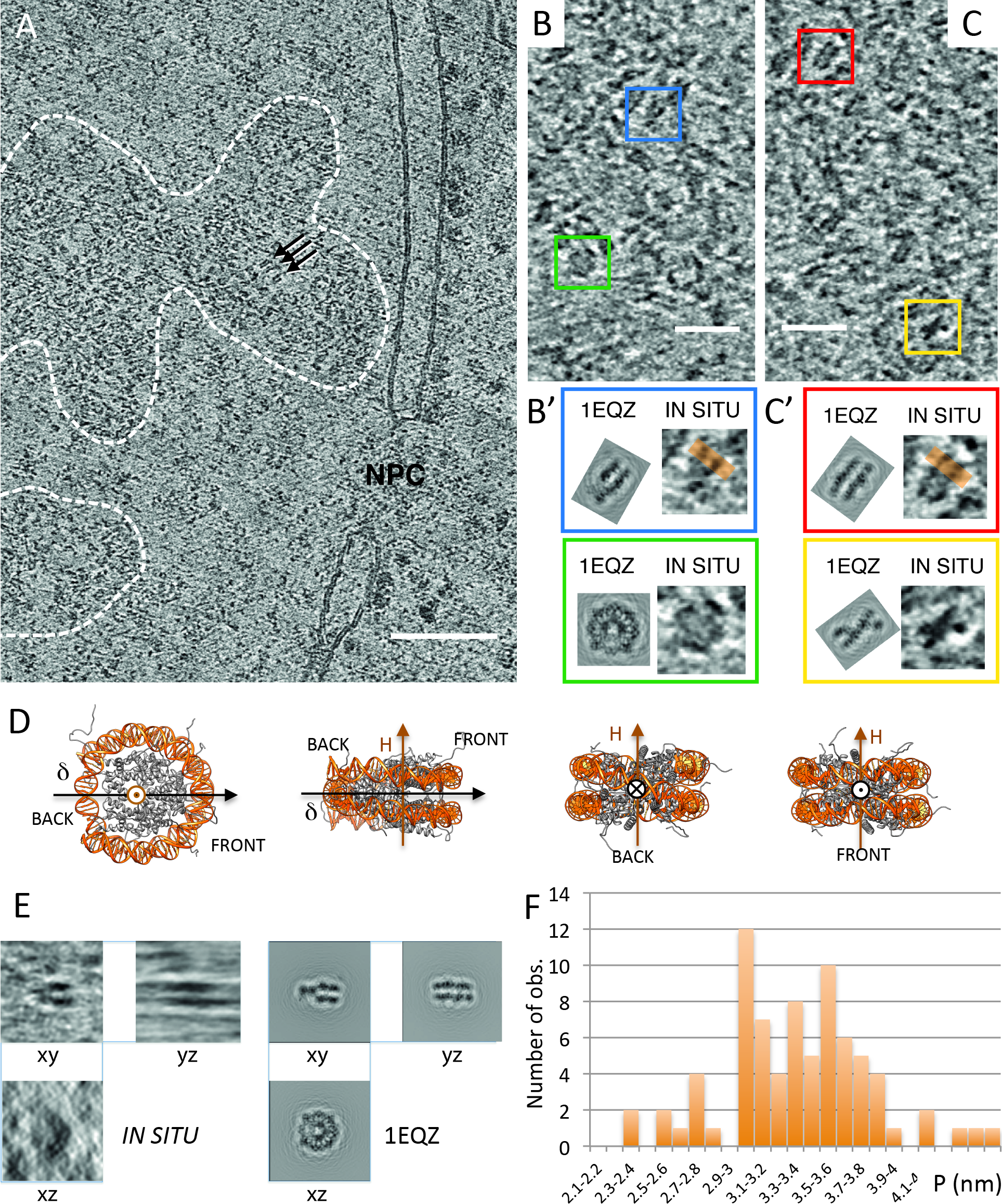
CETOVIS of interphase nuclei of embryonic fly brain. (A) A 5 nm thick slice through a tomographic reconstruction showing an area of nucleus (on the right) with a segment of nuclear envelope (NE) containing a nuclear pore complex (NPC). Dashed lines delineate two chromatin areas, in which individual nucleosomes are recognisable as dispersed 11 nm densities. (B, C) Enlargements of the chromatin areas show four characteristic patterns of 5 nm thick nucleosome sections, outlined with coloured square frames. Those are a round top view (green), and a family of side views, with a “v-shaped” section through the pseudo-dyad symmetry axis (blue), a striated front view (red) and an “x-shape” back view showing the entry/exit DNA regions (yellow). These patterns are compared with the corresponding CTF-modulated sections generated from the crystallographic structure of nucleosome (from PDB 1 EQZ). (D) Sketch of different orientations of the NCP (1EQZ) with its superhelical (H) and dyad (δ) axes. (E) Three perpendicular views of a nucleosome selected from the reconstructed volume are compared with the corresponding views of CTF-modulated crystallographic structure. The horizontal smearing visible in the YZ view is a missing wedge artefact. (F) Distribution of *P.*

#### Measurement of the distance P between DNA gyres: the nucleosome conformation in *situ* is variable and more open than the canonical crystallographic structure

On 2D images, wherever striated nucleosomes patterns were recognised (e.g. Figure 3D, D’, E, E’), *P* was systematically measured on intensity line profiles (Figure 3 D’, E’). Within HT29 and KE37 cell nuclei, *P* varies from 2.35 to 4.1 nm, with an average value of 3.13 ± 0.36 nm. The distributions of the *P*-values are shown in Figure 3F for the two types of cells. In both cases the distribution is normal (SW tests p-values respectively equal 0.399 and 0.168) and no significant difference was observed between the two cell lines (Student test p-value 0.518). For *Drosophila* embryonic brain cells *P* measurements were obtained from line profiles on computational slices of the reconstructions (orange lines in Figure 4B’C’). *P* also shows a normal distribution (SW test p-value 0.45) with an average value 3.37 nm (s.d. 0.45 nm) (Figure 4F).

As we did above for nucleosomes in solution, we compare these data with projections of the canonical crystallographic structure mimicking our imaging conditions (Supplementary Figure S5 and S6 for 2D and 3D data respectively). Relative to the canonical PDB structure, *in situ* measurements show a much broader distribution and a shift toward high *P*-values: 3.13 ± 0.36 nm *vs* 2.79 ± 0.14 nm for 2D data, and 3.37 ± 0.45 nm *vs* 2.98 ±0.24 for 3D ones. Nucleosomes observed *in situ* in frozen hydrated interphase nuclei on average significantly differ from the particle’s canonical structure. As we found for purified NCP in low salt solutions, they present a whole range of conformations and are more open than the canonical structure.

Note that the measured *P*-values depend on the imaging conditions (voltage and underfocus values), as analysed in SI (Supplementary Figure S8). CTF correction does not correct for this effect (3D data are CTF-corrected; for 2D data, we checked that CTF-correction does not change measured *P*-values, see Supplementary Figure S1). Since imaging conditions differ for 2D and tomographic data acquisitions, it is not possible to directly compare, nor simply normalise, the 2D and 3D datasets. Yet, the ratio of the mean *P*-value of nuclear nucleosomes to that of the corresponding PDB reference is the same for the two datasets (1.12 and 1.13 respectively). This suggests that the different nuclear nucleosome populations are similar whatever the considered cell type.

## DISCUSSION

There exists several reports of ultrastructural eukaryotic chromatin analyses *in situ*, both in interphase nuclei and mitotic chromosomes, based on either conventional or cryoEM approaches. Conventional EM of chemically fixed or freeze-substituted and stained cells long failed to evidence nucleosomes in cell nuclei, until the work of Kizilyaprak et al (50) that identified 11 nm fibres and electron dense particles in euchromatin regions. Using electron spectroscopic tomography Fussner et al (51) could distinguish phosphorus-rich DNA periphery and phosphorus-poor core histone octamer in some nucleosome particles, but could not resolve the DNA gyres. A recently developed staining methodology, ChromEMT, enabled access to the local conformation of the chromatin chain in interphase and mitotic chromosomes: in some places, individual particles were evidenced as single densities, yet with no conformational information (52). CEMOVIS approaches so far typically describe chromatin as a homogeneous grainy compartment (42-45), where nucleosomes were only detected as an 11-nm signal in FFTs (42,53). The first cryo-electron tomography studies went further, and nucleosome were visualised as ellipsoidal densities approximately 10 nm-wide and 6 nm-thick (53-55).

Here, the use of CEMOVIS and CETOVIS of interphase eukaryotic nuclei let us visualise, recognise and analyse nucleosomes in their native nuclear environment. For the first time the nucleosome can be analysed to a level of detail that allows its comparison with PDB structures and retrieve information about its conformation. Measuring the distance *P* between DNA gyres wrapped around the nucleosome, we detect the occurrence of multiple conformations, with *P*-values on average larger than that of the PDB structure, and with variations of this parameter of more than 0.5 nm around the average. This was found exploring very different cell types, from human cell lines to embryonic insect cells, suggesting that, in the interphase nucleus, open nucleosomes predominate.

We are thus also able to compare nuclear nucleosomes with nucleosomes in solution, and analyse the influence of a variety of factors. We solubilised here native purified NCP at physiologically relevant particles concentrations (20-50 %), and explored two different ionic environments. We show that the *P*-distance is also variable in solution, and exhibits a salt-dependant behaviour. Open conformations, with *P*-values on average higher than that of the canonical particle, characterise low salt solutions (15 mM), while increasing added salt concentration to 160 mM shifts the nucleosome to a more compact state, close to the canonical structure.

Despite their highly conserved structure, nucleosomes are already recognised as pleiomorphic and dynamical entities, with a multiplicity of states related to their variability in chemical composition (DNA sequence, histone variants, post translational modifications), their interaction with other molecules (H1 histones, remodelling factors, enzymes), as well as their dynamics (reviews in 13, 56). These conformational changes are so far essentially documented *in vitro* and *in silico*, or hypothesised by theoretical approaches. They may involve histone (mostly at the level of their tails) and/or DNA conformational changes. We will focus now mainly on the conformation of nucleosomal DNA (57,58).

CryoEM provides snapshots of the nucleosome population, and the obtained conformational diversity reflects both static and dynamic variability. Both nuclear nucleosomes *in situ*, and purified native particles are indeed chemically diverse, in terms of DNA sequence, histone variant content and post-translational modifications. High resolution crystallographic and cryoEM structures have so far shown no detectable effect of histones tail acetylation or DNA methylation (59-61) and only minor changes in the presence of the histone variants (H3.3 (62) or embryonic H2A.Z (63)), apart enhanced flexibility of DNA ends in CENP-A centromeric particles (64). Although DNA sequence plays a role in the particle stability it was not shown to result in major structural changes (see for example (65)). Altogether, it is therefore likely that our observations mostly reflect the dynamics of the particle, at least in solution, where the many nuclear factors (e.g. chromatin remodelers, etc…), that may further alter nucleosome conformation (66,67), are absent. Dynamical conformational transitions have been predicted by theoretical approaches and simulations (68-73), and probed by diverse experimental methodologies, from electron microscopy to AFM and single molecule manipulation and FRET (10,74-77). For example, the partial uncoiling of DNA, or “breathing” occurs spontaneously and involves breaking of histone-DNA contacts at the entry/exit sites of the DNA (8,9,78,79). Chiral transitions have also been found, under torsional stress (80,81).

We question the possible nature of the conformational changes evidenced in our study. On account of the salt dependence, we may hypothesise that the larger *P* values in low salt environment results from the electrostatic repulsion between the negatively charged DNA gyres, which would be screened at high salt. Indeed, the Debye length in low salt solution is in the range of *P* (3 nm at 10 mM), while it falls below 1 nm above 100 mM. The increase of DNA stiffness in low salt environment could also be involved.

Both in solution and *in situ*, it occurs along the direction of the DNA superhelical axis, perpendicular to the nucleosome plane. The variations of our measured *P*-values could describe different modes of out of plane nucleosome opening: (i) a global variation of the DNA superhelical pitch; or (ii) an edge opening/closing, or “gaping”, of the particle.

(i) An out of plane axial transition was first proposed by Marky and Manning (54), corresponding to an increase of the total superhelical pitch, as the result of elastic instability of DNA. Global out of plane motions in the direction of the superhelical axis were also found in molecular dynamics simulations (82), but with much smaller amplitudes (about 0.1 nm whereas about 0.5 nm here). Yet, the simulations were performed at the nanosecond time scale, while there could be DNA movements in the microsecond time scale and beyond. Note that extended nucleosome conformations, associated with an increase of the DNA superhelical pitch, have already been reported in two EM studies of dried and freeze-dried particles. A transition from an elongated oblate to a prolate, compact, ellipsoid has been reported with the increase of monovalent salt concentration from 10 to 150 mM (83); hyperacetylated nucleosomes at low ionic strength have been found extended (84). But the reported particles’ dimensions can hardly be reconciled with our cryoEM measurements, probably on account of dehydration associated with freeze-drying.
(ii) Later on, the gaping transition, corresponding to an edge opening of the nucleosome (like a gaping oyster), was proposed by Mozziconacci et al (69,70) as the result of stacking interactions to optimize chromatin condensation. Interestingly, a recent FRET analysis of synthetic 147 bp-601 particles in dilute solution has reported a change in the distance between DNA gyres on the front side of the particle, i.e. an edge opening characteristic of a gaping movement (85). It occurs through about 0.5 to 1 nm, an opening range in good agreement with our measurements. The transition is described as a slow switch between two discrete states: a compact state (with smaller distance between DNA gyres corresponding to the canonical crystallographic structure), and a gaped open state. The authors report a salt dependence: increasing the ionic strength increases the dwell time in compact conformation, and could also increase the compactness of the two states (85). Our *P* distribution in high salt solutions could be indicative of such a switch mechanism, with accumulation of compact canonical conformations (Figure 2D). However, in low salt solutions - as well as in interphase nuclei - we found normal *P* distributions, which rather suggests a continuum of conformations. This apparent discrepancy could be due to differences in methodology (the cryoEM approach traps intermediates of the slow gaping transition), but also to the nature of the particles (synthetic highly stable 601 nucleosomes *versus* native ones here). Indeed, the diversity of native nucleosomes could result in a range of slightly different conformations. In particular, DNA elasticity, which may play a role in a gaping transition, depends on its sequence. Differences in the experimental conditions (exact ionic environment and NCP concentration) could also play a role.

Whatsoever, the DNA conformational changes evidenced here are associated with changes in the histone octamer. These remain to be characterised and differ in the two situations considered above. At present, we have neither information about associated changes in the histone core, nor precise local understanding of DNA opening to tell apart the two possible modes of out of plane conformational changes. Sorting of back and front *P*-distances would be necessary to progress in that matter, but would require much larger sampling to obtain statistically reliable information. Note that it is possible that these two types of conformational changes may both take place, in solution and/or in interphase nuclei. More generally, further studies are necessary to understand this conformational change. Simulated cryoEM images obtained from gaped and extended nucleosomes atomic models should be compared to the experimental data. Hybrid approaches, such as the combination of FRET and cryoEM, have proved powerful in ribosome studies (86). The use of single particle cryoEM methods developed to access continuous conformational changes (87,88) would also be needed to characterise the conformational modes of the nucleosome.

We now question the relationship between the conformational changes evidenced in solution and those found in interphase nuclei.

Interestingly, open conformations that predominate *in situ* are favoured *in vitro* in “low salt” environment, while so-called “physiological” salt conditions (160 mM) favour compact canonical-like structures. Although counterintuitive, low added salt solutions may be more relevant to account for chromatin behaviour at high particle concentration. Indeed, the net negative charge of the NCP is about 150 (in the absence of post-translational modification and binding of additional basic proteins). In dilute solutions, the contribution of its counterions is negligible relative to the added ions. But, at a particle concentration of 25% (about 1.2 mM), the cation concentration to maintain electroneutrality reaches 180 mM. The NCP counterions then contribute significantly, leading to much higher total ionic concentrations.

We propose that the spontaneous opening occurring in low salt solution could account, at least in part, for the distribution of conformations evidenced *in situ.* However, the *P* distribution *in situ* is significantly higher and broader than in solution. This is not surprising, since nucleosomes are there not only chemically diverse but also interact with other nuclear components such as protein remodelers and enzymes. They can be expected to exist in a larger multiplicity of states, bound or not to various proteins, and in various local ionic environments, that remain to be explored and deciphered. Furthermore it is likely that our observations only describe part of conformational variability of nucleosomal DNA *in situ.* We did identify here only those particles that resemble closely the canonical structure. In particular, we did not search for more exotic particles that can be expected, such as hemisomes, hexasomes, reversomes (13, 89). To progress in this direction and access these states, CETOVIS, allowing us to follow the helical path of the DNA wrapped on single nucleosomes, is a very promising tool.

Spontaneous fluctuations and salt-dependant transitions evidenced *in vitro* could constitute a repertoire of configurations that specific regulatory processes can exploit. Local changes within the nuclear ionic environment (for example the increase in salt concentration induced by release of counterions upon binding of basic proteins), could serve as regulatory mechanism. Although mainly documented *in vitro*, for lack of a precise knowledge of local intracellular composition, this regards multiple levels of chromatin conformation: histone tails, that condense or extend (90), inter-particle interactions (91,92), order-disorder transitions as well as the local tuning of chromatin concentrations (17,92,93), and, at large scale, coil-globule transitions of chromosome domains (94,95).

As regards the possible functional implications of nucleosome conformational changes, attention has been so far mainly turned toward breathing movements. Indeed, certain histone variants and post-translational modifications are associated to transient unwrapping of DNA, while there is no evidence that out of plane opening, such as gaping, is involved in biological processes. In contrast with breathing, gaping could stand a way to favour access to nucleosomal DNA without disrupting DNA-histone contacts. According to the theoretical model of Mozziconacci et al (69,70), the gaping transition would be triggered by nucleosome stacking to promote condensation, and isolated gaped nucleosomes would not exist. But this is contradicted by the FRET observations in dilute solutions (85). Here, *in vitro*, we found neither effect of particle concentration nor of their relative orientation (parallel or antiparallel, see Figure 1B, F) on the NCP opening. However, we analysed too small a range of conditions (concentrations ranging from 25 to 32% at low salt, and around 45% at high salt) where, in all cases, nucleosomes stack into columns. Preliminary results (not shown) on isolated nucleosomes in dilute solution suggest that open conformations also occur in the absence of stacking, in good agreement with Ngo et al (85). In interphase nuclei *P*-values were recorded both on stacked and not stacked particles (see Figure 3 D, E) and we did not sort the two situations.

The relationship between nucleosome opening and the chain topology has also to be envisaged. Interestingly enough, open conformations are found with isolated purified particles as well as with linked ones in the chromatin fibre. Mozziconacci et al (69,70) suggested that gaping would be accompanied by twist and would thus also drive supercoiling. Yet, their model relies on the topological properties of a 30nm helical fibre, which is not the case in the more disordered state found in the interphase nucleus (52, 96). Nonetheless, curvature of the DNA superhelical axis associated with gaping lowers the energetic cost of positive supercoiling (97), and gaping could therefore be used to regulate locally DNA topology and contribute to the capability of chromatin to accommodate high levels of supercoiling, (“topological buffer”(98)). The interplay between the chain topology, the stacking/unstacking and the conformation of the particle remains to be explored.

To question the physiological significance of nucleosome opening it will also be necessary to target *in situ* measurements. Correlative light-electron microscopy (CLEM) approaches should be used to map nucleosomes and evidence possible conformational differences between particles found in hetero- and euchromatin regions, telomeres or centromeres for example.

## Acknowledgements

We thank J. J. Lacapère and L. Issop (Faculté X. Bichat, Paris) and A.M. Tassin (I2BC, Gif-sur-Yvette) for providing HT29 and KE37 cells respectively. We thank G. Pehau-Arnaudet for his support at the federative FEG-cryo-electron microscope (Institut Pasteur, Paris) and J. Dubochet (Univ. Lausanne) for access and support at the LAU (Uni. Lausanne, Switzerland). We thank H. Gnaegi (Diatome) for providing diamond knives for tomographic experiments and M. Kunz (Univ. Frankfurt) for help with tilt series reconstruction. This work was funded by the program ANR-06-PCVI-0029. This work was supported by “Investissements d’Avenir” LabEx PALM (ANR-10-LABX-0039-PALM). M. Eltsov and D. Grewe were supported by the German Research Foundation (DFG) grant EL 861/1-1.

